# Cigarette Smoke Exposure Impairs β-Cell Function Through Activation of Oxidative Stress and Ceramide Accumulation

**DOI:** 10.1101/761007

**Authors:** Xin Tong, Zunaira Chaudry, Chih-Chun Lee, Robert N. Bone, Sukrati Kanojia, Judith Maddatu, Paul Sohn, Morgan A. Robertson, Irina Petrache, Carmella Evans-Molina, Tatsuyoshi Kono

**Author notes:** Address correspondence and requests for reprints to: Carmella Evans-Molina, MD, PhD or Tatsuyoshi Kono, PhD, Indiana University School of Medicine, 635 Barnhill Drive, MS 2031A, Indianapolis, IN 46202, or Irina Petrache, MD, 1400 Jackson St, Denver, CO 80806. These authors contributed equally to this work (Xin Tong, Zunaira Chaudry).

## Abstract

**Objectives:** Epidemiological studies indicate that first- and second-hand cigarette smoke (CS) exposure are important risk factors for the development of type 2 diabetes (T2D). Additionally, elevated diabetes risk has been reported to occur within a short period of time after smoking cessation, and health risks associated with smoking are increased when combined with obesity. At present, the mechanisms underlying these associations remain incompletely understood. The objective of this study was to test the impact of CS exposure on pancreatic β-cell function using rodent and *in vitro* models.

**Methods:** Beginning at 8 weeks of age, C57BL/6J mice were concurrently fed high fat-diet (HFD) and exposed to CS for 11 weeks, followed by an additional 11 weeks of smoking cessation with continued HFD exposure. Glucose tolerance testing was performed during CS exposure and during the cessation period. Cultured β-cells (INS-1) and primary islets were exposed *ex vivo* to CS extract (CSE), and β-cell function and viability were tested. Since CS increases ceramide in lungs cells and these bioactive sphingolipids have been implicated in pancreatic β-cell dysfunction in diabetes, islet and β-cell sphingolipid levels were measured in islets from CS-exposed mice and in CSE-treated islets and INS-1 cells using liquid chromatography-tandem mass spectrometry.

**Results:** Compared to HFD-fed ambient air-exposed mice, HFD-fed and CS-exposed mice had reduced weight gain and better glucose tolerance during the active smoking period. Following smoking cessation, CS-mice exhibited rapid weight gain and a significantly greater increase in glucose intolerance compared to non-smoking control mice. CS-exposed mice had higher serum proinsulin/insulin ratios, indicative of β-cell dysfunction, significantly lower β-cell mass (p=0.02), and reduced β-cell proliferation (p=0.006), and increased islet ceramide accumulation. *Ex vivo* exposure of isolated islets to CSE was sufficient to increase islet ceramide accumulation, reduce *insulin* gene expression and glucose-stimulated insulin secretion, and increase β-cell oxidative and ER stress. Treatment with the antioxidant N-acetylcysteine, markedly attenuated the effects of CSE on ceramide levels, restored β-cell function and survival, and increased cyclin D2 expression, while also reducing activation of β-cell ER and oxidative stress.

**Conclusions:** Our results indicate that CS exposure inhibits insulin production, processing, and secretion and reduced β-cell viability and proliferation. These effects were linked to increased β-cell oxidative and ER stress and ceramide accumulation. Mice fed HFD continued to experience detrimental effects of CS exposure even during smoking cessation. Elucidation of mechanisms by which CS exposure impairs β-cell function in synergy with obesity will help design therapeutic and preventive interventions for both active and former smokers.

## 1. INTRODUCTION

Worldwide, diabetes and cigarette smoking are highly prevalent public health concerns. Type 2 diabetes (T2D) impacts over 422 million individuals, and more than 1.1 billion individuals actively smoke [1; 2]. More strikingly, nearly 1.9 billion individuals are exposed second hand to cigarette smoke (CS) [3]. While a number of genetic risk variants for T2D have been identified through genome wide association studies, lifestyle factors such as nutrition, physical activity, and potential chemical and environmental exposures are thought to strongly influence diabetes susceptibility. Epidemiological studies indicate that CS exposure is an important risk factor for the development of T2D, and this diabetes risk remains heightened even after smoking cessation [4; 5]. Thus, first and second hand smoke exposure represent important modifiable risk factors for the development of T2D in a potentially large pool of at-risk individuals [6; 7].

Human physiologic studies suggest that CS exposure impairs peripheral insulin sensitivity and pancreatic β-cell function, despite a link to reduced body weight [8-10]. In addition, individual cohort studies showed that within 1-5 years after smoking cessation, subjects with a previous smoking history had a 22-91% higher incidence of developing T2D [11; 12], and this increased risk was correlated to weight gain after cessation of tobacco use [13]. At present, the mechanisms underlying impaired glucose homeostasis in response to cigarette smoke exposure, especially a full understanding of the effects of smoking on pancreatic β-cell function, remain unknown.

To address this knowledge gap, we developed a rodent model of CS exposure coupled with high-fat diet to mimic the combined effects of cigarette smoking and diet-induced obesity. To provide specific insight into mechanisms whereby cigarette smoke exposure impacts β-cell health, analysis of this rodent model was combined with an *in vitro* model where cultured β-cells and primary rodent islets were exposed to CS extract (CSE). Exposure to CSE, derived from smoke condensate followed by removal of particulate matter, has been extensively used in *ex vivo* models [14-18]. Previous studies have reported a significant increase in ceramide in the lung and other tissues after experimental exposure to CS [18-20], so we also tested whether CS-exposure influenced β-cell ceramide content. In our *in vivo* model, the combination of CS exposure and HFD caused rapid weight gain in mice and β-cell ceramide accumulation during the immediate cessation period. Smoking-induced ceramide accumulation in β-cells was correlated with impaired β-cell function, reduced β-cell mass and proliferation, and impaired insulin secretion and processing. These effects were largely recapitulated in our *in vitro* model of CS-exposed β-cells and islets, where we observed increased ceramide levels, reduced insulin secretion and reduced β- cell survival. The antioxidant NAC reversed the *in vitro* effects of CS exposure and prevented excessive ceramide accumulation, suggesting a primary role for oxidative stress in smoking-induced β-cell dysfunction.

## 2. MATERIALS AND METHODS

### 2.1. Animals and metabolic studies

Male C57BL/6J mice were obtained from Jackson Laboratories (Bar Harbor, ME) at 7 weeks of age and maintained under protocols approved by the Indiana University School of Medicine Institutional Animal Care and Use Committee in accordance with guidelines from the Association for Assessment and Accreditation of Laboratory Animal Care. Mice were fed a high fat-diet (HFD) containing 45% of kilocalories from fat (D12451, Research Diets Laboratories, New Brunswick, NJ), beginning at 8 weeks of age and were maintained on a standard light-dark cycle with *ad libitum* access to HFD and water. In parallel with HFD initiation, mice were exposed to CS using 3R4F research grade cigarettes (Kentucky Tobacco Research and Development Center, University of Kentucky, Lexington, KY), with 11% mainstream and 89% side-stream smoke or ambient air control for 5 hours a day, 5 days a week for a total of 11 weeks, using the Teague 10E whole body exposure apparatus (Figure 1A) [21]. Intraperitoneal glucose tolerance tests (GTT) were performed after 6 hrs of fasting followed by the administration of glucose at a dose of 2 g/kg total body weight. Insulin tolerance tests (ITT) were performed after 3 hrs of fasting and administration of recombinant human insulin from Boehringer Ingelheim Vetmedica (Duluth, GA) at a dose of 0.75 IU/kg total body weight. Glucose levels were measured using the AlphaTRAK glucometer (Abbott Laboratories, Abbott Park, IL). Serum insulin and proinsulin levels were measured using ELISAs from Mercodia (Salem, NC) and ALPCO Diagnostics (Salem, NH), respectively. Dual X-ray Absorptiometry (DEXA) analysis was performed to estimate body composition using the Lunar PIXImus II (GE Medical Systems) as previously described.

**Figure 1.**
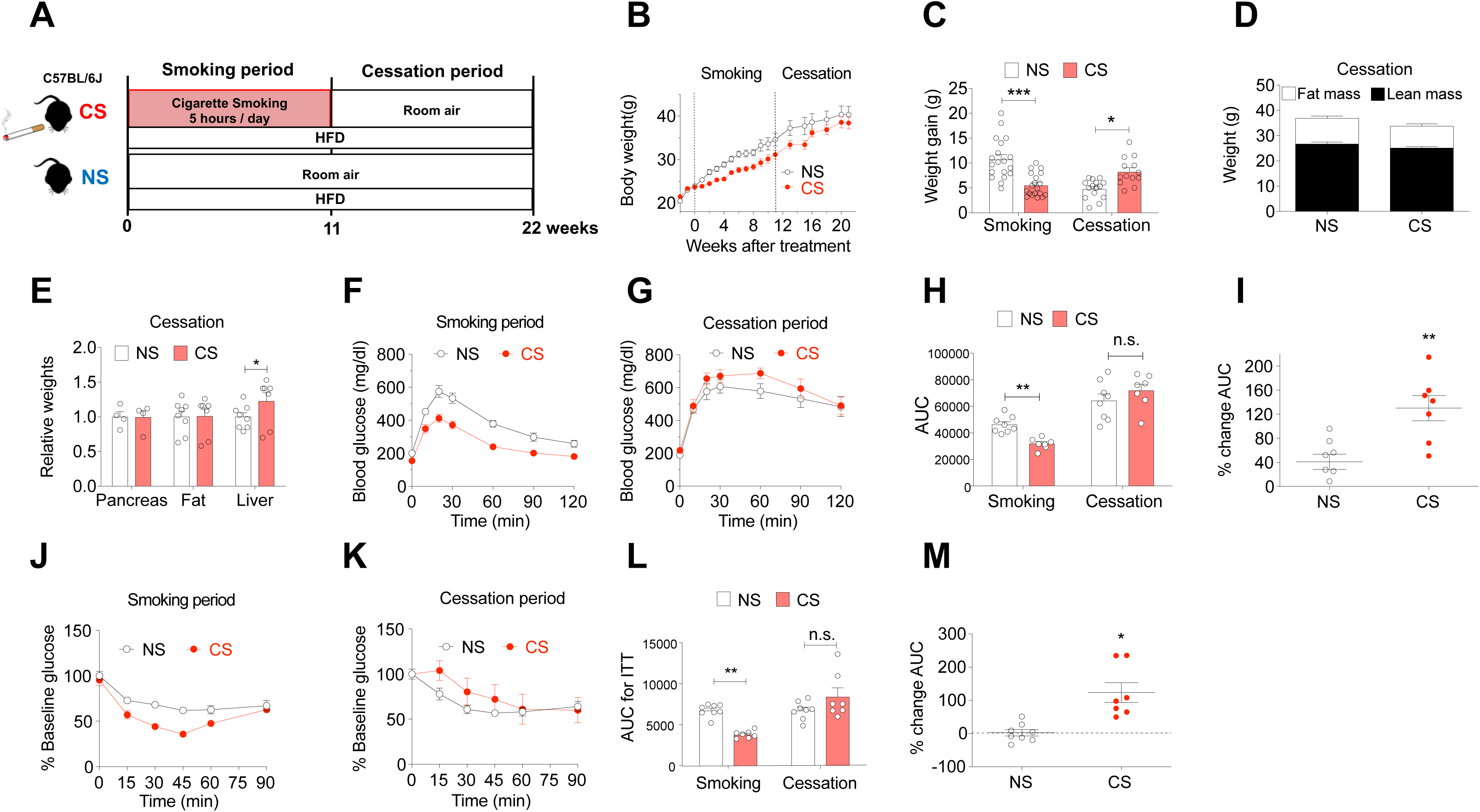
CS exposure increased weight gain after smoking cessation and led to impaired glucose tolerance in HFD-fed C57BL/6J mice. (A) Schematic of study design. Non-smoking (NS) and Cigarette Smoke (CS) groups were fed a high- fat diet (HFD) containing 45% of kilocalories from fat for 22 wks. CS mice were placed in a smoking chamber for 5 hrs/day for 5 days per week for the first 11 weeks. NS mice were exposed to room air. After 11 weeks, both NS and CS groups continued for an additional 11 weeks with standard housing conditions. (B) Results of weekly body weight measurements for 0-22 wks of the study. (C) Body weight gain during the smoking and cessation periods. (D-E) Body lean/fat mass and weights of whole pancreas, liver, and epididymal fat pads were compared between the groups at the end of the study. (F-G, J-K) Glucose tolerance tests (GTT) and insulin tolerance tests (ITT) were performed during the smoking period (week 9-11) and the cession period (week 20-22). (H, L) Area Under Curve (AUC) for GTT and ITT data was quantified. (I, M) Percentage (%) change of AUC during the cessation period (week 20-22 minus week 9-11 / week 9-11) was analyzed for each group. Data for individual animals are indicated by the open circles; n=8-15. Results are displayed as the means ± S.E.M; *p<0.05, **p<0.01, ***p<0.001 compared to non- smoking group (NS) or indicated comparisons.

### 2.2. Immunohistochemistry and immunofluorescence

Pancreata were rapidly removed following euthanasia and fixed overnight using Z-fix buffered zinc formalin fixative (Anatech Ltd., Battle Creek, MI), followed by paraffin embedding and longitudinal sectioning at 5 μm intervals as previously described [22]. β-cell proliferation was assessed as detailed in previous publications [23]. β-cell mass was estimated for each animal by determining the average β-cell surface area multiplied by the pancreatic weight as previously described [23; 24].

### 2.3. Glucose-stimulated insulin secretion (GSIS), Quantitative real-time PCR (qRT-PCR) and immunoblot

Mouse pancreatic islets were isolated by collagenase digestion and INS-1 832/13 rat insulinoma cells were cultured as previously described [25]. Static GSIS, qRT-PCR, and immunoblot analysis were performed using previously published methods [26]. Islet perifusion based secretory profiles was measured using the Biorep Perifusion System (Biorep, Miami Lakes, FL). Twenty-four hours after isolation, 50 handpicked islets were loaded into each perifusion chamber; islets were perifused with Krebs buffer containing 2.8 mmol/L glucose for 20 min, followed by 16.7 mmol/L glucose for 40 min at a rate of 120 μL/min. Secreted insulin was measured using ELISA (Mercodia); results were normalized to total DNA content. The following primary antibodies were employed at the indicated dilutions: anti-total caspase-3 rabbit antibody, which detected both cleaved and total caspase-3 (1:1000, Cell Signaling, Danvers, MA); anti-PC1/3 rabbit antibody (1:1000, Cell Signaling); anti-Cyclin D2 rabbit antibody (1:1000, Santa Cruz Biotechnology, Santa Cruz, CA), anti-Actin mouse antibody (1:10000, MP Bioscience, Santa Ana, CA).

### 2.4. Cigarettes smoke extract (CSE) preparation and cell-based studies

Aqueous CSE was obtained from research-grade cigarettes (3R4F) purchased from the Kentucky Tobacco Research and Development Center (University of Kentucky, Lexington, KY) as previously described [18]. In brief, air control (AC) or CSE stock (100%) extract were prepared by bubbling ambient air or smoke from two cigarettes into 20 ml of PBS at a rate of 1 cigarette/min, followed by pH adjustment to 7.4, and 0.2 µm filtration. Cell and islet treatments were performed using CSE or AC extract concentrations ranging from 1% to 10% (vol:vol). To adjust the variation of potencies of the CSE from different lots, we performed titration tests for cell viability measuring ATP levels with CellTiter-Glo Luminescent Cell Viability Assay (Promega, Madison, WI). We determined a lethal concentration defined as 50% death after 24h treatment (LC50), and used this concentration for CSE treatment. For propidium iodide (PI) staining, INS-1 cells were dissociated with trypsin-EDTA, followed by 70% ethanol fixation. Cells were then washed with PBS and incubated at 37°C for 40 min in 50 μg/ml PI solution. Imaging cytometry analysis was performed using the CellometerTM K2 (Nexcelom Bioscience, Lawrence, MA). CellTiter-Glo Luminescent Cell Viability Assay (Promega, Madison, WI) and Earlytox live/dead assay kit (Molecular Devices, San Jose, CA) were used for studies of INS-1 cells treated with CSE or AC extract with or without 5 mM N-Acetyl Cysteine (NAC) from Invitrogen; Thermo Fisher Scientific, Inc (Waltham, MA) and tauroursodeoxycholic acid (TUDCA) from Cayman Chemical (Ann Arbor, MI) coincubation for 24 h.

### 2.5. Analysis of sphingolipids by LC-MS/MS

To profile sphingolipids in isolated mouse islets or INS-1 cells, lipid extraction and liquid chromatography-tandem mass spectrometry (LC-MS/MS) were performed according to detailed methods published previously [19]. The MS system was Sciex 6500 QTRAP mass spectrometer interfaced with a Shimadzu Nexiera X2 UHPLC system. The sphingolipids were ionized via electrospray ionization (ESI) with detection via multiple reaction monitoring (MRM). Analysis of sphingoid bases and the molecular species of ceramides employed ESI in positive ions with MRM analysis. C17-sphingosine and N-C17-ceramide were used as the internal standards.

### 2.6. Statistical analysis

Differences between groups were analyzed for significance using unpaired Student’s t test or one-way analysis of variance with Tukey-Kramer post hoc analysis. Results are displayed as the means ± S.E.M. Data were analyzed using GraphPad Prism software (GraphPad Software, La Jolla, CA, USA) and a p value < 0.05 was taken to indicate the presence of a significant difference between experimental groups.

## 3. RESULTS

### 3.1. CS exposure followed by cessation increased weight gain and impaired glucose tolerance in mice

C57BL/6J mice were exposed to an initial phase of 11 weeks of CS exposure (for 5 hours/day; 5 days per week) combined with 45% HFD, which closely mimics a high-fat Western diet [27]. This was followed by smoking cessation and continued HFD exposure for an additional 11 weeks (Fig. 1A). Control mice (NS) were maintained under normal ambient air control conditions combined with 45% HFD for the entire 22-week treatment period.

During the active smoking period, CS-exposed mice had reduced weight gain compared to the NS group, but their rate of weight gain increased during the smoking cessation period (Fig. 1B-C), such that by the end of the 22-week experiment, total body weights (Fig. 1B), lean mass, and fat mass (Fig. 1D) were similar between the groups. Whole pancreas, liver, and epididymal fat weights were analyzed, and only liver weights were significantly higher in the CS-exposed group at week 22 (Fig. 1D-E).

Concordant with the observed patterns of total body weight, glucose tolerance in CS-exposed mice at 11 weeks was better than the HFD-mice maintained under control air conditions. However, GTT responses in CS mice dramatically worsened during the smoking cessation period (Fig. 1F-H). When analyzed as % change in the area under the curve (AUC) of the GTT response from 11 to 22 weeks, CS-exposed mice exhibited a significantly higher % change (Fig. 1I). In addition, the insulin sensitivity of the CS-exposed group was higher during the smoking period, followed by marked worsening during the smoking cessation period (Fig.1J-L). Similar to the glucose tolerance trend, the CS group animals had a significantly higher % change in the AUC of the ITT response from 11 to 22 weeks (Fig.1M).

### 3.2. CS reduced insulin processing, β-cell mass, and β-cell proliferation during the smoking cessation period in C57BL/6J mice

To define the origins of the failed adaptive response to HFD seen during the smoking cessation period, we analyzed insulin processing capacity, β-cell mass and β-cell proliferation in CS-exposed and NS mice at 22 weeks. Elevations in the serum proinsulin/insulin (PI/I) ratio are observed in humans with both T1D and T2D and reflect a reduced ability to complete processing of proinsulin to mature insulin arising within the context of increased β-cell stress [4]. The PI/I ratio in serum was increased in the CS-exposed group compared to NS controls (Fig. 2A-C). Analysis of β-cell mass revealed a significant reduction (3.11 mg vs. 2.25 mg, p=0.02) in CS-exposed mice (Fig. 2D). Consistent with this finding, β- cell proliferation, which was determined by proliferating cell nuclear antigen (PCNA) and insulin co-staining, was reduced in CS-exposed mice compared to control mice (3.98 % vs. 1.86 %, p=0.006) (Fig. 2E).

**Figure 2.**
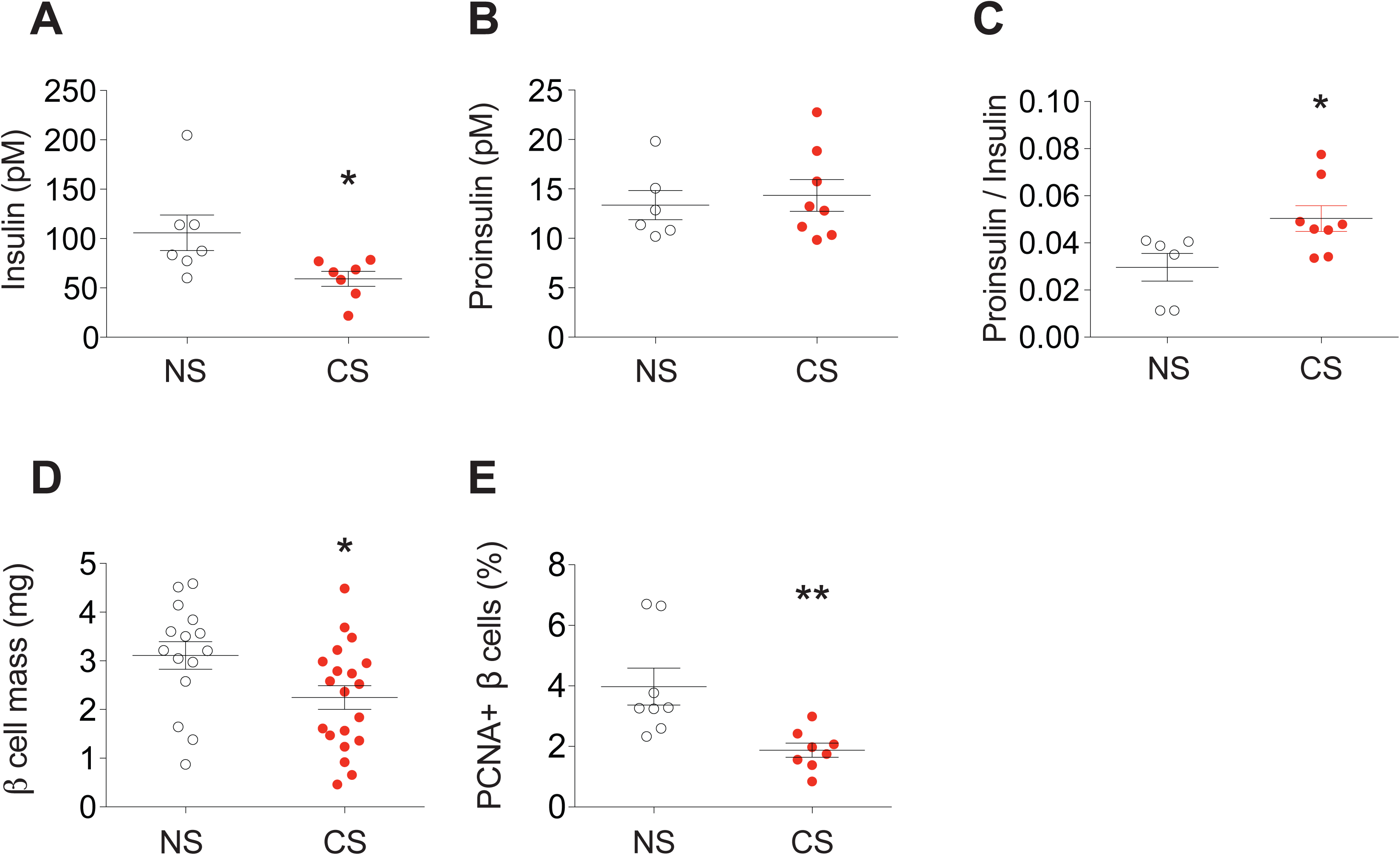
CS reduced insulin processing, β-cell mass, and β-cell proliferation during the smoking cessation period in C57BL/6J mice. Following the study design depicted in Fig. 1A, the capacity for insulin production was evaluated in NS and CS mice at the study endpoint of 22 wks. (A-C) Random blood was collected and serum levels of insulin, proinsulin, and the ratios of proinsulin to insulin were determined by ELISA. (D-E) The mass of insulin positive cells and the number of proliferating β-cells within pancreata of study mice were quantified in paraffin tissue sections by immunostaining for insulin and the proliferative marker PCNA. Data for individual mice in each group are indicated by the circles. Results are displayed as the means ± S.E.M; *p<0.05, **p<0.01 compared to non-smoking (NS) group.

### 3.3. CS exposure increased ceramide accumulation in pancreatic islets during the smoking cessation period

Next, using tandem mass spectrometry, we measured ceramide levels in isolated islets from mice exposed to CS before and following smoking cessation. At week 11, CS exposure was associated with modest reductions in ceramide compared to control mice, mostly in palmitoyl (C16:0)- and behenic (C22:0)- ceramide species. In contrast, CS-exposed mice exhibited marked increases in ceramide in islets isolated at 16-weeks, affecting multiple species including stearic (C18:0)-, arachidic (C20:0)-, docosanoid-, and the saturated and monounsaturated lignoceric (C24:0 and C24:1)- ceramides (Fig. 3B).

**Figure 3.**
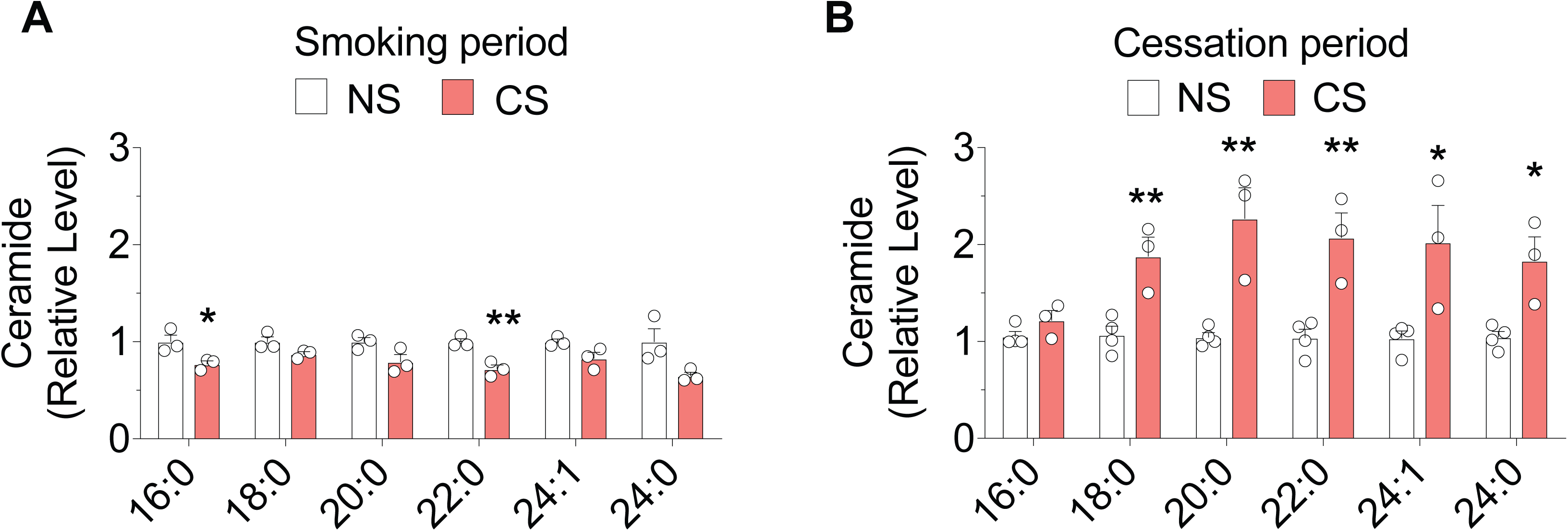
CS exposure increased ceramide accumulation in pancreatic islets following smoking cessation. The study design of Fig 1A was executed and pancreatic islets were isolated during the last wk of the smoking period (A) and 5 wks after smoking cessation (B). The concentrations of 6 different ceramide species were determined by LC-MS/MS and are shown as relative levels. Each sample has a total 200 islets pooled from two mice. The replicates for each experimental group are indicated by the open circles and the number of replicated animals in each group was n=3-4. Results are displayed as the means ± S.E.M; *p<0.05, **p<0.01, ***p<0.001, compared to non-smoking group (NS) group.

### 3.4. CSE treatment increased ceramides and related metabolites in INS-1 cells and isolated mouse islets

Islets could be directly affected by soluble components of CS that cross the alveolar-capillary lung barrier and are distributed to tissues *via* circulation. Alternatively, they may be injured by mediators released by other CS-activated cells in a paracrine or autocrine manner [28; 29]. To determine if CS exposure was by itself sufficient to cause islet dysfunction and ceramide accumulation, we exposed cultured β-cells and mouse islets to CSE *ex vivo*. In both models, CSE exposure for 24 hours resulted in a marked increases in multiple ceramide species. Compared to control-treated cells, CSE increased ceramides in INS-1 cells from ∼1.4 fold (palmitoyl- and stearic-ceramides) to more than 1.7 fold (behenic- and lignoceric- ceramides) (Fig. 4A). Ceramides were increased to an even greater extent in CSE-exposed islets, with most species being increased by more than 3-fold compared to control conditions (Fig. 4A-D). Mass spectrometric measurements of other related sphingolipid metabolites revealed increases in both precursors of ceramide in the *de novo* sphingolipid synthesis pathway (DH sphingosine and DH-ceramide (Fig. 4B-D) and, in the case of isolated islets, sphingosine (Fig. 4D), which is an immediate downstream ceramide metabolite, produced by *via* ceramidase.

**Figure 4.**
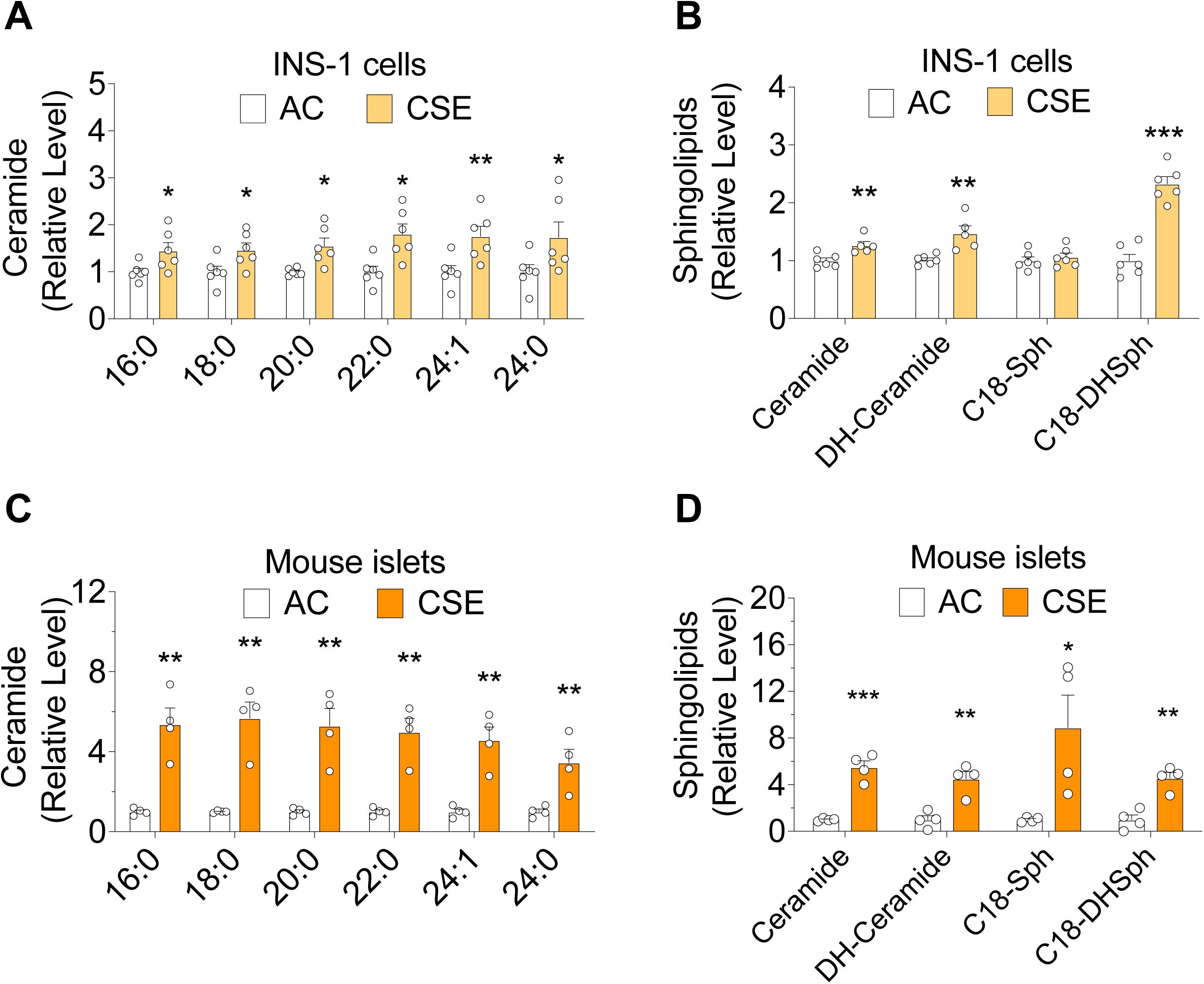
CSE treatment altered ceramide synthesis in INS-1 cells and isolated mouse islets. Cells and isolated mouse islets were incubated for 24 hrs with cigarette smoke extract (CSE) or air control (AC) extract that were prepared by bubbling ambient air or smoke at a rate of 1 cigarette/min. (A and C) The concentrations of 6 different ceramides species were determined by LC-MS/MS. (B and D) Summary of LC-MS/MS results for total ceramide, dihydro(DH)-ceramide, C (18:0)-spingosince, and C(18:0) dihydro(DH)-sphingosine. The replicates of cell samples and islets samples are indicated by open circles. For cells, n=5-6. For islets, each sample was prepared from a total 200 islets pooled from two mice, with a total 8 individual mice per group. Results are displayed as the means ± S.E.M; *p<0.05, **p<0.01, ***p<0.001, compared to AC group.

### 3.5. Exposure to CSE reduced insulin secretion and increased activation of β- cell ER stress and oxidative stress signaling pathways

Results from our *in vivo* model showed that serum insulin levels were lower in mice exposed to CS. To test directly the impact of CSE on insulin secretion and production, insulin expression was measured and glucose stimulated insulin secretion (GSIS) assays were performed in INS-1 cells exposed to CSE for 24 hrs. GSIS was significantly reduced (284.5 vs 115.2 μU/ml/mg protein, p=0.005) in CSE-exposed INS-1 cells versus control cells (Fig. 5A). In addition, both mature and pre-insulin mRNA levels were reduced by 70.1% and 68.7%, respectively in CSE-exposed cells (Fig. 5B-D), indicating a defect in active insulin gene transcription as well as a reduction in total levels of mature insulin mRNA.

**Figure 5.**
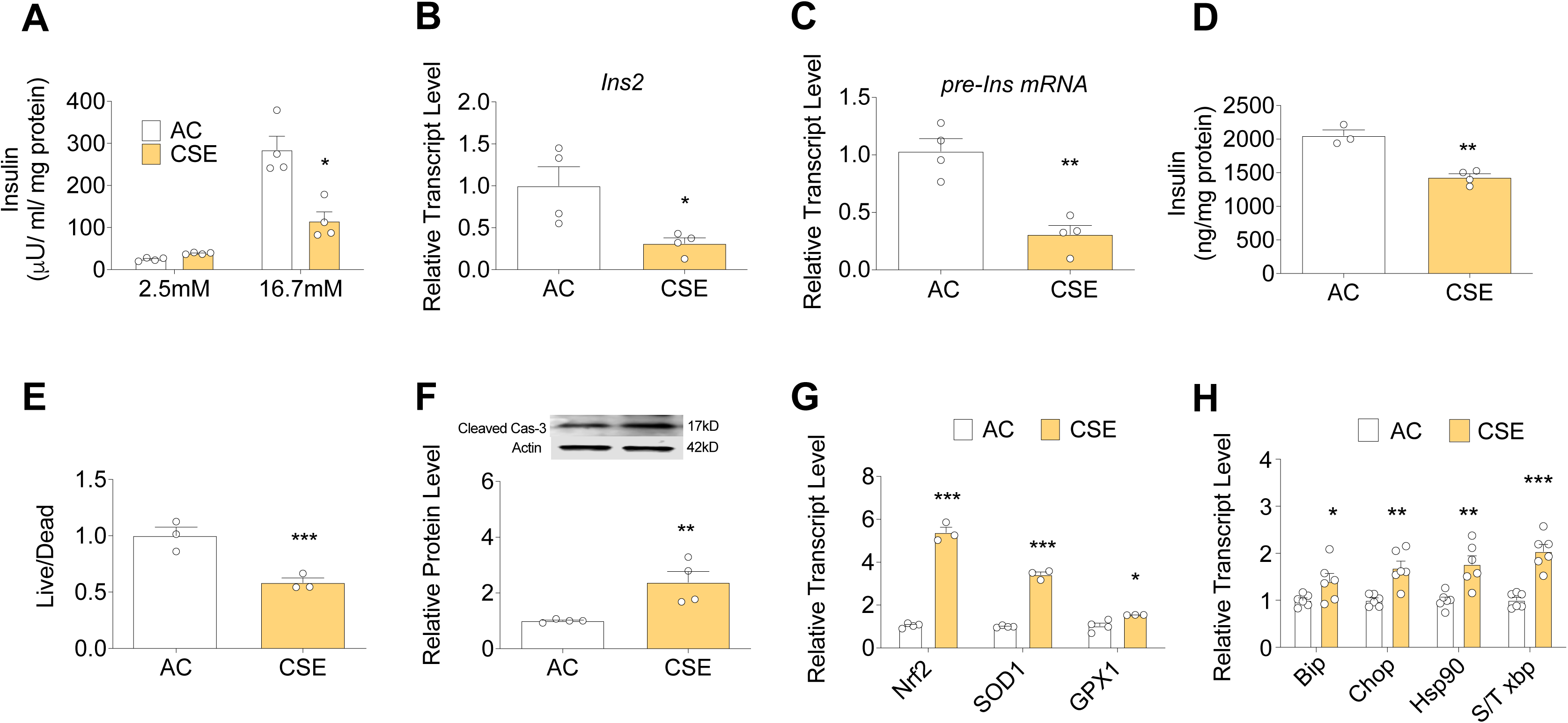
Exposure to CSE reduced insulin secretion and increased activation of β-cell ER stress and oxidative stress. Cells were incubated for 24 hrs with cigarette smoke extract (CSE) or air control (AC) extracts as in Fig. 4. (A) Glucose-stimulated insulin secretion into the supernatant was measured by ELISA and results were normalized to cellular protein levels. (B-C) RNA was isolated from treated INS-1 cells and insulin transcript levels were determined by qRT-PCR. (D) Cellular insulin levels were determined by ELISA with normalization of values to total protein levels. (E) Cell viability was determined by live/dead assay, and results are presented as the ratio of live to dead cells. (F) Immunoblotting was performed using caspase-3 and actin antibodies in control and CSE treated cells. Results were normalized with actin expression levels. (G) Cells were incubated for 6 hrs with CSE or AC extracts and gene expression of the oxidative stress markers *Nrf2, Sod1* and *Gpx1* were determined by qRT-PCR. Results were normalized with *Actb* expression levels. (H) Cells were incubated for 24 hrs with CSE or AC extracts. Expression of the ER stress marker genes, *Bip, Chop, Hsp90*, and spliced *Xbp-1* ratio were analyzed as described in (G). The number of replicate samples for each analysis is indicated by the open circles; n=3-6. Results are displayed as the means ± S.E.M; *p<0.05, **p<0.01, ***p<0.001 compared to AC group.

Since both elevations in the proinsulin/insulin ratio, reduced insulin expression, and ceramides have been linked with activation of β-cell oxidative stress and ER stress [18], we next quantified the expression of genes involved in these processes. Results from this analysis indicated a global increase in the expression of genes involved in oxidative stress (*Nrf2, Sod1, and Gpx1)* and ER stress signaling *(Bip, CHOP, Hsp30,* and spliced/total *Xbp-1*) (Fig. 5G-H). Consistent with these findings, live/dead analysis showed that β-cell survival was reduced by ∼41.5% following CSE exposure, while immunoblot analysis revealed a significant increase in cleaved caspase-3 expression in CSE-exposed β-cells (Fig.5E-F). Taken together, these results suggest that CSE exposure directly induced ceramide accumulation in islets and β-cells, which was linked with reduced insulin secretion, impaired β-cell survival, and β-cell ER and oxidative stress.

### 3.6. Antioxidant treatment prevented CSE-induced loss of β-cell function and viability

To test whether approaches aimed at reversing either oxidative or ER stress could prevent CSE-induced beta cells dysfunction, we utilized the antioxidant n-acetylcysteine (NAC: 5 mM) and the chemical chaperone tauroursodeoxycholic acid (TUDCA: 100 µM). The antioxidant effect of NAC was confirmed by its inhibitory effect on CSE-induced upregulation of *Nrf2,* a master transcriptional regulator of the cellular antioxidant response (Fig. 6A). As expected, TUDCA treatment prevented CSE-induced upregulation of ER stress markers (Fig. 6B). Interestingly, ER stress was suppressed not only by TUDCA but also by NAC treatment, suggesting that CSE-induced ER stress may be linked to induction of oxidative stress.

**Figure 6.**
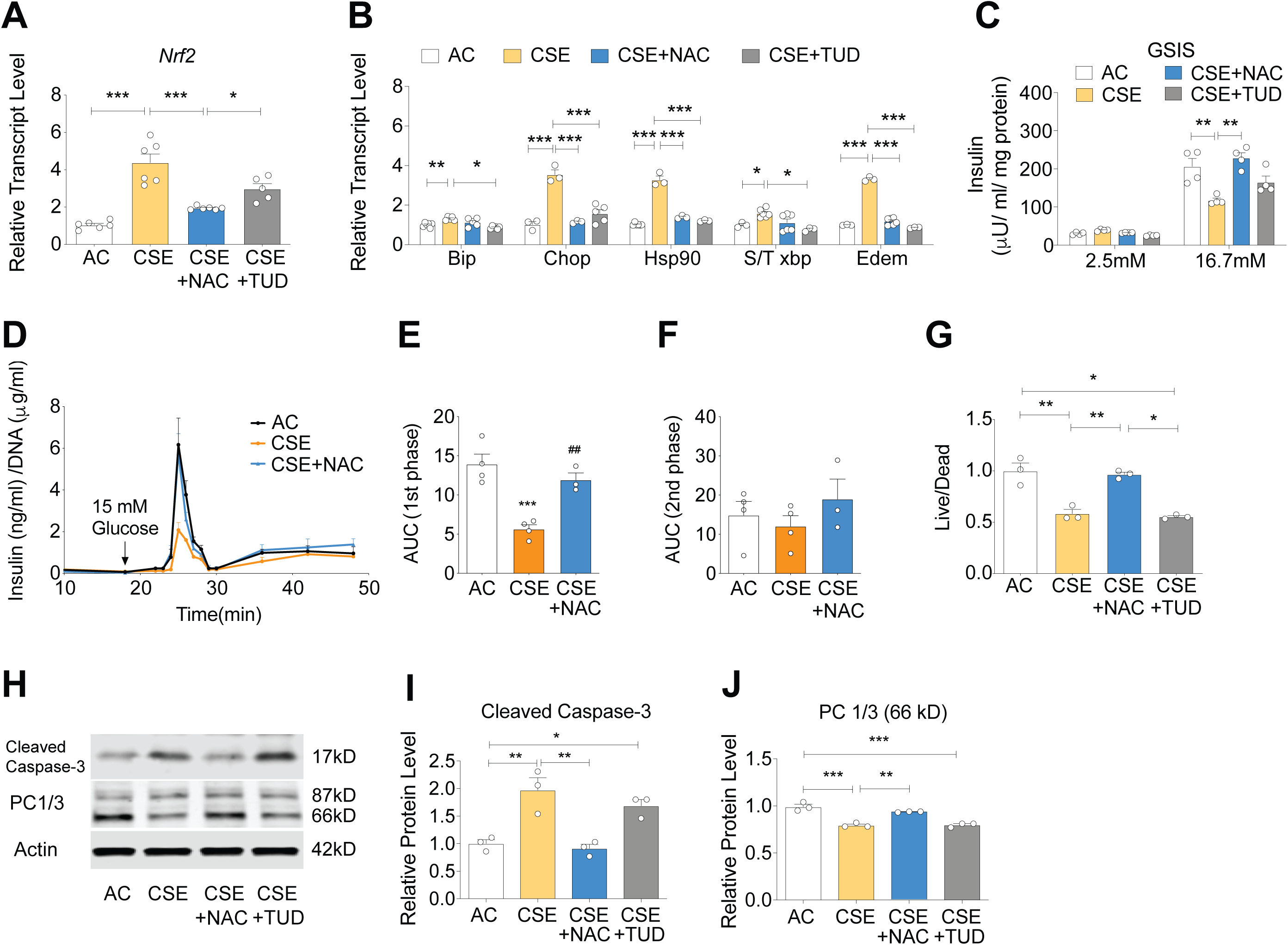
Antioxidant treatment prevented CSE-induced loss of β-cell function and viability. INS-1 cells were co-incubated with N-acetylcysteine (NAC: 5 mM) or the chemical chaperone TUDCA (TUD: 100 µM) and cigarette smoke extract (CSE) for 24 h prior to assessment of indices of β-cell function. (A-B) RNA was isolated from treated INS-1 cells and gene expression of the oxidative stress marker *Nrf2* and ER stress markers, *Bip, Chop, Hsp90*, and spliced *Xbp-1* ratios were determined by qRT-PCR. (C) Glucose-stimulated insulin secretion into the supernatant was measured by ELISA, and results were normalized to cellular protein levels. (D-F) Mouse islets were treated with CSE for 24 hrs and loaded into perifusion chambers and then perfused with Krebs buffer containing 2.8 mmol/L glucose for 20 min, followed by 16.7 mmol/L glucose for 40 min at a rate of 120 μL/min. Secreted insulin was measured using ELISA, and the results were normalized to total DNA content. (G-I) Total protein was isolated from treated cells and immunoblotting was performed using antibodies recognizing caspase-3, Cyclin D2, proprotein convertase and actin. Results were normalized to actin levels. (J) Cell viability was determined by a live/dead assay, and results expressed as the ratio of live to dead cells. The numbers of replicates for each measurement are indicated by the open circles, n=3-4. Results are displayed as the means ± S.E.M; *p<0.05, **p<0.01, ***p<0.001 for indicated comparisons.

Treatment with NAC, but not with TUDCA attenuated the effects of CSE on GSIS in INS-1 cells, suggesting a major role for oxidative stress in CSE-induced impairments in β-cell function (Fig. 6C). To further characterize the impact of NAC on phasic insulin secretion in CSE-treated mouse islets, perifusion experiments were performed. Here, CSE-treatment led to a marked reduction in first phase insulin release, while insulin secretion was significantly restored upon NAC treatment (Fig. 6D-F). Similar to effects observed on insulin secretion, NAC treatment inhibited CSE-induced changes in β-cell viability as measured by live/dead assays (Fig. 6G). Moreover, NAC but not TUDCA reduced caspase-3 cleavage (Fig. 6H-I).

Finally, considering the critical role that prohormone convertases play in proinsulin processing, we measured expression levels of PC1/3 in our *in vitro* model. Our results showed robust (21%) down-regulation of the active form [30] (66 kD) of PC1/3 by CSE. In addition, NAC, but not TUDCA treatment, was able to restore PC1/3 expression in CSE-treated INS-1 cells (Fig.6J).

### 3.7. Antioxidant treatment prevented the effect of CSE on sphingolipid accumulation in isolated mouse islets

Since our data suggested that antioxidant treatment, rather than selective prevention of ER stress, was necessary to mitigate the effects of CSE on β-cell function, we next tested whether NAC also inhibited CSE-induced ceramide accmumulation using LC-MS/MS. Isolated mouse islets treated with NAC (5 mM, 24 hrs) while exposed to CSE showed significantly reduced ceramide accumulation (Fig. 7A-B) across all ceramide species measured, by more than 55% (P<0.05), as well as other sphingolipid metabolites proximal to ceramide de novo synthesis (DH-sphingosine and DH ceramide) and immediately downstream of ceramide (sphingosine) compared to untreated CSE-exposed islets. The increase of another pro-apoptotic ceramide metabolite, hydroxy-ceramide [31] induced by CSE was markedly decreased by NAC (Fig. 7B). These data indicated that more than a half of the excessive sphingolipid accumulation induced by CSE was oxidative stress-dependent.

**Figure 7.**
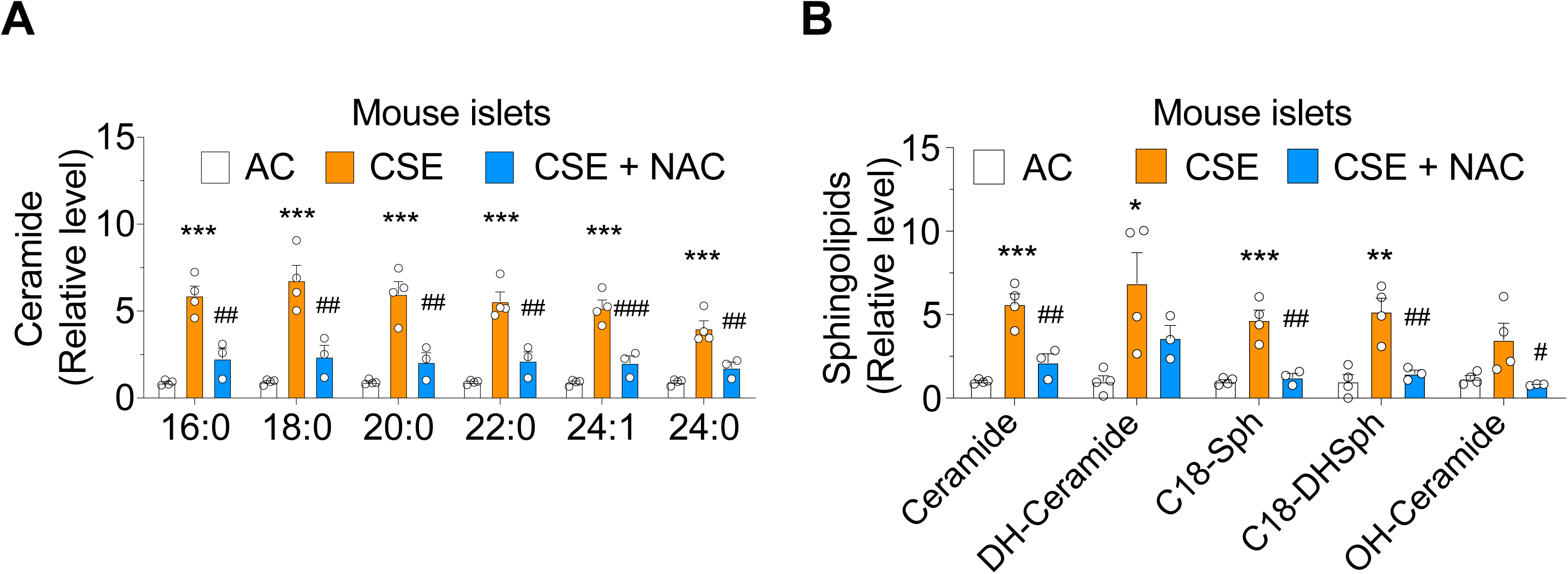
Antioxidant treatment prevented the effect of CSE on lipid accumulation in isolated mouse islets. Mouse islets were co-incubated with N- acetylcysteine (NAC: 5 mM) and cigarette smoke extract (CSE) for 24 h prior to analysis of lipid levels. (A-B) Summary of LC-MS/MS results for total ceramide, dihydro(DH)-ceramide C (18:0)-spingosine, C (18:0) dihydro(DH)- sphingosine measurement, and hydroxy-ceramide. Each sample was prepared from total 200 islets pooled from two mice, total 6-8 individual mice per group. The numbers of replicates for each experiment are indicated by the open circles, n=3-4. Results are displayed as the means ± S.E.M; *p<0.05, **p<0.01, ***p<0.001 compared to AC, ^#^p<0.05, ^##^p<0.01, ^###^p<0.001 compared to CSE group.

### 3.8. CS down-regulated CyclinD2 and impaired cell proliferation in pancreatic islets

To further understand how CS exposure reduced β-cell mass and proliferation, we interrogated cell cycle status in INS-1 cells after 24h exposure to CSE. CSE reduced cell proliferation, which was characterized by an increased percentage of cells in the G0/G1 phase (39.2 % vs. 47.7 %; p<0.05) and a decreased percentage of cells in the G2/M phase (45.3 % vs. 37.4 %; p<0.05) (Fig. 8A). qRT-PCR analysis of a broad panel of cell cycle regulators revealed decreased *cyclin D2* mRNA levels, while *cyclin D1, and E1* mRNA levels were increased in CSE-treated cells (Fig.8B). Immunoblot analysis in INS-1 cells treated with CSE for 24h, showed a significant decrease in cyclin D2 protein expression (Fig.8C). Consistent with this result, CSE reduced *cyclin D2* mRNA levels in isolated mouse islets (Fig.8D). To determine the relevance of these findings in our *in vivo* smoking model, immunostaining of cyclin D2 was performed in pancreatic sections collected from CS and NS-exposed mice at the 22-week timepoint. CS-exposed mice exhibited significant reductions in cyclin D2 expression within islets, supporting the notion that β-cell proliferation was reduced *via* loss of this critical cell cycle regulator (Fig. 8E).

**Figure. 8.**
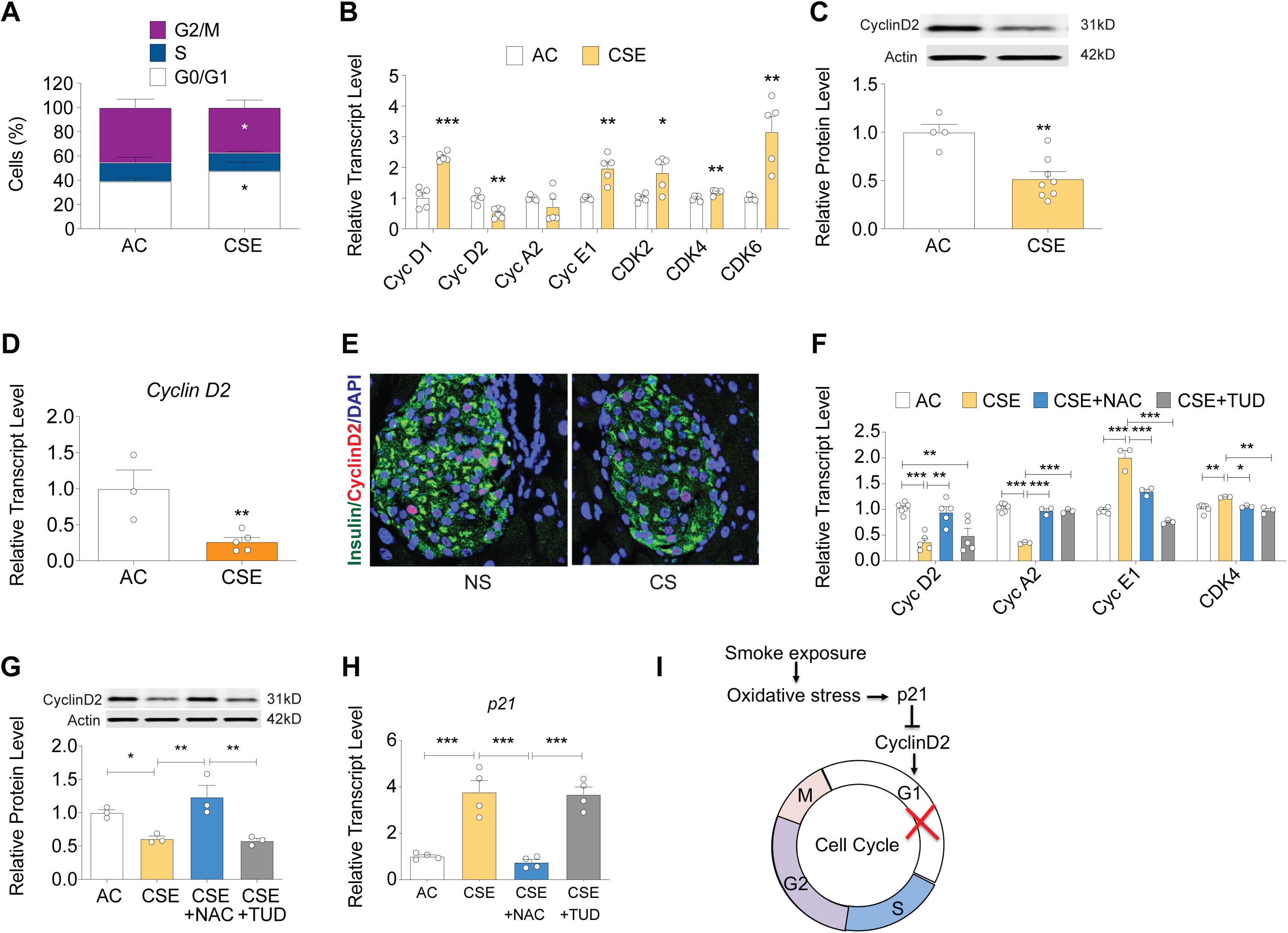
CS impaired β-cell cell proliferation through cyclin D2 loss. (A) INS- 1 cells were incubated with AC or CSE for 24 hrs and stained with propidium iodide (PI), and cell cycle analysis was performed using flow cytometry. (B) Total RNA was isolated from treated INS-1 cells, and gene expression of a panel of cyclin genes was performed (Cyclin D1 (*Ccnd1*), D2 (*Ccnd2*), D3 (*Ccnd3*), A2 (*Ccna2*), E1 (*Ccne1*), G1 (*Ccng1*), *Cdk2, Cdk4, Cdk6* were determined by qRT-PCR. (C) Total protein was isolated from treated cells and immunoblotting was performed using antibodies against Cyclin D2 and actin. Results were normalized to actin levels. (D) Mouse islets were treated with 5% CSE for 24h and qRT-PCR of *CyclinD2* was performed. (E) Immunostaining images of paraffin sections of pancreata from C57BL/6J mouse treated with the combination of CS and HFD for 22 wks. (F-H) INS-1 cells were co-incubated with N-acetylcysteine (NAC: 5 mM) or the chemical chaperone TUDCA (TUD: 100 µM) and CSE for 24 h prior to assessment of gene or protein expression. RNA was isolated from treated INS-1 cells and gene expression of potent cyclin-dependent kinase inhibitor *p21(Cdkn1a)* and cyclin genes (*Ccnd2, Ccna2, Ccne1, Cdk4*) were determined by qRT-PCR (F and H). Cyclin D2 protein level was determined by immunoblotting (G); (I) Hypothesized pathway for oxidative stress-dependent cell cycle arrest in β-cell. The numbers of replicates for each experiment are indicated by the open circles. Results are displayed as the means ± S.E.M; *p<0.05, **p<0.01, ***p<0.001 compared to AC treatment.

Finally, we tested a role for oxidative stress in cell-cycle regulation in our CSE-model. Treatment with NAC significantly attenuated the changes in cyclin D2 expression induced by CSE exposure (Fig. 8F-G), suggesting an important role for oxidative stress in cell cycle regulation. Using the same treatment conditions, we determined the expression levels of *cyclin-dependent kinase inhibitor p21*, which is a known negative regulator of cyclin D2 [32]. Realtime qPCR analysis revealed a significant increase in *p21* expression by CSE, which was prevented by treatment with NAC (Fig. 8H). Together, these results suggested that β-cell proliferation was reduced due to oxidative stress through upregulation of *p21* coupled with a reduction in cyclin D2 expression (Fig.8I).

## 4. DISCUSSION

Despite an abundance of epidemiological data that links tobacco smoking with an increased risk of diabetes, little is known regarding the mechanisms underlying these associations. We aimed to address this knowledge gap by using a murine *in vivo* model of chronic CS exposure combined with HFD complemented with an *in vitro* model of acute CS exposure using INS-1 cells and isolated islets exposed to soluble components of CS. To the best of our knowledge, this is the first study to investigate the mechanisms by which CS impacts β-cell health and function.

Notably, we included in our experimental design assessments of direct effects of CS measured during a relatively chronic period of CS exposure, as well as effects of CS that were sustained following an equal period of CS cessation. Extrapolating lifespans of mice to humans, these timeframes have been linked with metabolic dysfunction and diabetes risk. In persons with and without diabetes, cigarette smoking is independently associated with higher hemoglobin A1c (HbA1c) concentrations [33], and diabetes risk increases in a dose-dependent manner with the number of cigarettes smoked per day [34]. Interestingly, the risk of diabetes is heightened in individuals who have recently quit smoking, and this risk has been found to correlate with weight gain following cessation. Therefore, the mouse model used in our studies is highly relevant to exploring the mechanisms of human metabolic dysregulation that links CS, obesity, and hyperglycemia.

As previously reported in CS-exposed mice fed regular chow [18], mice displayed reduced total body weight gain during CS exposure, even when fed HFD. This observation is consistent with known effects of CS to suppress appetite and increase energy expenditure [20]. Glucose and insulin tolerance largely followed weight patterns, and mice exposed to CS were more glucose tolerant and more insulin sensitive compared to HFD-fed non-exposed mice. However, following cessation of CS exposure, mice exhibited an accelerated body weight gain, with associated worsened glucose and insulin tolerance. Thus, our model recapitulated the elevation of T2D risk after smoking cessation in terms of both weight gain and glucose tolerance. However, despite equivalent body weights between CS exposed and control mice at the end of the 22-week observation period, CS-exposed mice had reduced β-cell mass and proliferation, reduced serum insulin levels, and increased proinsulin/insulin ratios, suggesting that the chronic CS exposure reduced the capacity for β-cell adaptation in the setting of continued HFD exposure. These mice were also found to have increased ceramide levels within their islets.

The lung, which is directly exposed via inhalation to the toxic effects of tobacco smoking, exhibits a multitude of cellular responses to chronic CS exposure that include oxidative stress, inflammation, lung cell injury that results in cell death and tissue remodeling, as well as neoplasia [35]. In particular, in pulmonary emphysema, which is primarily caused by chronic CS, excessive ceramide production has been a major contributor to the apoptotic loss of lung endothelial cells that participate in gas exchange [19]. CS exposure in mice increased ceramide production in other organs, including cardiomyocytes, leading to disrupted cardiomyocyte mitochondrial function [20], and the skeletal muscle [36]. In addition to a local production of ceramides in various tissues, typically as a result of CS-induced oxidative stress [37] or mitochondrial dysfunction[38], systemic ceramides are increased during CS and can cause tissue damage. There are numerous studies of how CS affects serum lipid profiles in general – most focusing on cholesterol and associated lipoproteins- and how these are only partially reversible upon smoking cessation [39]. Regarding ceramides in particular, CS alone can result in the release of ceramide-rich endothelial exosomes and microparticles into the systemic circulation [40]. Profiling of metabolites in the serum of smokers and ex-smokers indicated an accelerated production of ceramide in those affected with chronic lung disease [41]. Further, a study of former smokers that also evaluated the impact of obesity and high fat diet on plasma lipids, suggested that these were associated with elevated levels of serum ceramides [42]. Thus, islets could be indirectly affected by CS-induced increases in circulating ceramides or by other soluble components of CS that cross the alveolar-capillary lung barrier and are distributed to tissues via the circulation [20; 43]. Alternatively, they could be injured by mediators released by other CS-activated cells in a paracrine manner [28; 29]. A third alternative is that CS activates a program of increased ceramide production directly within the islet or β-cell.

To determine the direct effects of CS on β-cells and isolated mouse islets, we exposed them *ex vivo* with CSE and noted that it markedly decreased β-cell proliferation, viability, insulin biosynthesis, maturation, and secretion. In addition, ceramide levels were increased within both CS-exposed islets and INS-1 β-cells, suggesting that CS directly impacts ceramide synthesis pathways in the β-cell. Although we did not directly measure the expression and the activity of the enzymes involved in the complex ceramide metabolism, the elevation of DH ceramide and DH sphingosine suggested that at least the *de novo* pathway of ceramide synthesis may account for ceramide accumulation. The elevation in sphingosine suggest that ceramidase is also activated to hydrolyze the excessive ceramide and/or there is an inhibition in the metabolism of sphingosine to its downstream metabolite sphingosine-1 phosphate (S1P). Of these sphingolipid metabolites, DH ceramide and sphingosine are known to have a role in cellular autophagy [38] [44], whereas the biological effects of DH-sphingosine and hydroxy ceramide have been less well defined, the latter having been shown to be pro-apoptotic.

Multiple reports, recently reviewed by Boslem *et al* indicate that the accumulation of ceramide and other sphingolipids have a major impact driving pancreatic β-cell dysfunction in both T1D and T2D [45]. Sphingolipid metabolites such as ceramide, sphingosine-1-phosphate, and sphingomyelin has been linked with activation of oxidative and ER stress [45], β-cell apoptosis [46], disruption of vesicle-trafficking [47], and inhibition of insulin gene expression [48]. Concurrently, many of these pathways are also reported to be involved in smoking related toxicity in other cell types [18; 19]. Indeed, gene expression of both oxidative and ER stress markers were upregulated in our *in vitro* model of smoke exposure, indicating a critical role for these stress pathways in smoking related β-cell failure. Although prior studies suggested the importance of ER stress in ceramide-induced apoptosis and impaired β-cell function [45; 49], our results demonstrated that rescue of ER stress by the chemical chaperone TUDCA did not mitigate the overall effects of CSE. In contrast, antioxidant treatment with NAC markedly reduced CSE-induced apoptosis, normalized expression of PC1/3 and Cyclin D2, and restored first phase insulin secretion. Interestingly, NAC reduced CSE-induced ceramide accumulation as well as ER stress, suggesting that oxidative stress from CSE could play a determining role in ceramide increases and could be responsible for β-cell apoptosis and dysfunction. A similar effect of NAC on CSE-induced cell death was reported previously in lung endothelial and epithelial cells [18; 50]; however, our study is the first to suggest that NAC may have beneficial effects on the β-cell in smoking models. Tobacco smoke contains over 4000 noxious chemicals as well as free radicals and reactive oxygen species (ROS), with many reported to show potential for biological oxidative damage due to smoking [51; 52]. In addition, it has also been reported that ceramide accumulation can further increase oxidative stress [37], *via* several mechanisms, including direct effects on the mitochondrial membrane [53]. Thus, oxidative damage could be induced by free radicals and ROS, which can be amplified when combined with ceramide accumulation. Although further experimentation is needed, our data suggest a potential mutual, self-amplified interaction of ceramide and ROS in CS induced- β-cell dysfunction.

The role of individual ceramide species in disease pathogenesis in general and in diabetes in particular is far from being fully elucidated. However, our data is consistent with other reports in models of diabetes, where ceramides with palmitoyl, behenic, and unsaturated lignoceroyl fatty acids have been reported elevated in streptozotocin models of diabetes in rats [54]. Further, in a study of mice fed HFD, mice deficient in the enzyme that metabolized sphingosine to S1P (SPhK1) were more likely to develop diabetes associated with enhanced β-cell apoptosis, further augmented by palmitic fatty acids, whereas S1P had a pro-survival effect [55]. Although we did not measure S1P levels, the further elevations in sphingosine and palmitoyl ceramide in CS-exposed mice and islet cells are consistent with these results in HFD models.

There are limitations to our study that should be considered. First, our primary focus was to elucidate pathways associated with smoking induced pancreatic β- cell dysfunction. Human physiological data has shown that smoking also impacts insulin sensitivity. Indeed, we observed evidence of worsened insulin resistance in our *in vivo* studies, but we did not explore mechanisms associated with this finding. Our intent was to create an *in vivo* model that fully recapitulated human responses to smoking. However, we did not observe worsened glucose tolerance during the smoke exposure period, likely due to the complexity of nicotine signaling on appetite and energy expenditure. We overcame this limitation by coupling our *in vivo* studies with an *in vitro* model of CSE-treatment in β-cells and islets. Finally, our study utilized male mice only, so we cannot extrapolate these results to female animals.

Notwithstanding these limitations, our data identify several critical mechanisms through which smoking impacts pancreatic β-cell function and survival. We found that smoking led to ceramide accumulation that was linked with activation of oxidative and endoplasmic reticulum stress. At a mechanistic level, these changes were associated with reduced insulin production, impaired insulin processing and reduced insulin secretion as well as reduced β-cell viability and proliferation. Taken together, these data provide novel insights into the mechanisms whereby cigarette smoking increases the risk of T2D during both the smoking period and during the immediate cessation period.

## 5. CONCLUSION

In summary, we conclude that CS exposure elevates the risk of impaired glucose homeostasis through β-cell dysfunction caused by oxidative stress and ceramide accumulation.

## List of abbreviations

AUC: Area under the curve
BiP/GRP78/HSPA5: Binding immunoglobulin protein, Glucose regulated protein 78kD, Heat-Shock 70kD protein 5
CHOP: CCAAT/enhancer-binding protein homologous protein
CS: Cigarette smoke
CSE: Cigarette smoke extract
GPX1: Glutathione Peroxidase 1
HFD: High-fat diet
Hsp30: Heat-Shock protein 30 kD
GTT: Glucose tolerance test
ITT: Insulin tolerance test
LC-MS/MS: Liquid chromatography-tandem mass spectrometry
NAC: n-acetylcysteine
NS: Non-smoking control
Nrf2: Nuclear factor erythroid 2–related factor 2
PC1/3: Proprotein convertase 1/3
p21: cyclin-dependent kinase inhibitor p21
SOD1: Superoxide dismutase 1
T2D: Type 2 diabetes
Xbp-1: X-box binding protein 1

## Author Contributions

X.T. and Z.C. design of the study, data analysis and interpretation, collection and assembly of data, and manuscript draft writing. C. L., R.N.B., S.K., J.M., P.S., and M.A.R. participated in data collection and critical revision of the manuscript. I.P. contributed to data analysis, provided critical reagents, and provided critical revision of the manuscript. C.E.-M.: directed funding acquisition, study conception and design, directed manuscript writing, and gave final approval of the manuscript. T.K. directed funding acquisition, conception and design of the study, data analysis and interpretation, collection and assembly of data, and manuscript writing/editing. T.K., I.P., and C.E.-M. are the guarantors of this work, had full access to all of the study data, and take responsibility for the integrity and accuracy of the data.

## Acknowledgement

The authors would like to thank Dr. Donalyn Scheuner (Indiana Bioscience Research Institute) for her helpful advice and discussions. The authors would also like to thank Dr. Evgeny Berdyshev (National Jewish Health) for his assistance with ceramide analysis. We thank Dr. Kathleen Heidler, and Jacob Saliba (Indiana University), for their technical support for *in vivo* smoke exposure experiments. We also thank Kara Orr and Karishma Randhave (Indiana University) for their technical assistance.

