## SupplementalFig.S1S2 for "Cigarette Smoke Exposure Impairs β-Cell Function Through Activation of Oxidative Stress and Ceramide Accumulation"

Supplemental table S1. Rat Primers used in current study

| Full name | Gene name | Sequence (5’ to 3’) |
| --- | --- | --- |
| Insulin | Ins2 | F: TCTTCTACACACCCATGTCCC  R: GGTGCAGCACTGATCCAC |
| Insulin exon 2–intron 2 | Ins2E2-I2 | F: GGGGAGCGTGGCTTCTTCTA  R: GGGGACAGAATTCAGTGGCA |
| Nuclear factor erythroid 2-related factor 2 | Nrf2 | F: TGGATGAAGAGACCGGAGAA  R: CAAGGCATCTTGTTTGGGAATG |
| Superoxide dismutase | SOD1 | F: CGTCATTCACTTCGAGCAGA  R: GCTCGCCTTCAGTTAATCCT |
| Glutathione peroxidase 1 | Gpx1 | F: GGCAAGGTGCTGCTCAT  R: CGCTTCTGCAGATCATTCATTTC |
| Heat Shock Protein Family A Member 5 (BiP) | Grp78 | F: CCACCAGGATGCAGACATTG  R: AGGGCCTCCACTTCCATAGA |
| The C/EBP Homologous Protein (CHOP) | Ddit3 | F: CCACCACACCTGAAAGCAGAA  R: AGGTGAAAGGCAGGGACTCA |
| Heat shock protein 90 | Hsp90 | F: ACACCACAGAAGACACCACAGATG  R: GGGAGAGGGAGGCTTGGG |
| X-box binding protein 1 | Total-Xbp1 | F: AGCACTCAGACTACGTGCGCCTC  R: CCAGAATGCCCAAAAGGATATCAG |
|  | Spliced-Xbp1 | F: CTGAGTCCGCAGCAGGT  R: TGTCAGAGTCCATGGGAAGA |
| ER degradation-enhancing alpha-mannosidase-like protein 1 (Edem) | Edem1 | F: CTACCTGCGAAGAGGCCG  R: GTTCATGAGCTGCCCACTGA |
| CyclinA2 | Ccna2 | F: TCCTTGCTTTTGACTTGGCT  R: ATGACTCAGGCCAGCTCTGT |
| CyclinD1 | Ccnd1 | F: GCGTACCCTGACACCAATCT  R: CACAACTTCTCGGCAGTCAA |
| CyclinD2 (Mouse and Rat) | Ccnd2 | F: GCTATGGAGCTGCTGTGCT  R: CCAAGAAACGGTCCAGGTAA |
| CyclinE1 | Ccne1 | F: GCTTCTAGACCTGTGCGTCC  R: CTTTCTTTGCTTGGGCTTTG |
| Cyclin dependent kinase 2 | CDK2 | F: GTTGACGGGAGAAGTTGTGG  R: TGATGAGGGGAAGAGGAATG |
| Cyclin dependent kinase 4 | CDK4 | F: TATGAACCCGTGGCTGAAAT  R: CCTTGATGTCCCGATCAGTT |
| Cyclin dependent kinase 6 | CDK6 | F: GCCTATGGGAAGGTGTTCAA  R: GGGCTCTGGAACTTTATCCA |
| Cyclin dependent kinase inhibitor 1A (p21) | Cdkn1a | F: CCTGGTGATGTCCGACCTG  R: CCATGAGCGCATCGCAATC |
| Actin | Actb | F: AGGTCATCACTATTGGCAACGA  R: CACTTCATGATGGAATTGAATGTAGTT |

Supplemental table S2. Antibodies used in current study

| Targeted protein | Species | Manufacturer | Dilution |
| --- | --- | --- | --- |
| PC1/3 | Rabbit | Cell signaling | 1:500 |
| CyclinD2 | Rabbit | Cell signaling | 1:1000 |
| Caspase 3 | Rabbit | Cell signaling | 1:1000 |
| Insulin | Guinea pig | Invitrogen | 1:250 |
| PCNA | Rabbit | Santa Cruz | 1:100 |
| Actin | Mouse | MP Biomedicals | 1:10000 |
